# Heterologous production, purification and crystallization of 24C-sterol methyltransferase from *Candida albicans*

**DOI:** 10.1101/2025.02.05.636639

**Authors:** Nokwanda S. Mpontshane, Michael Fairhead, Diederik J. Opperman, Rodney Hart, Valmary van Breda, Frank von Delft, Lizbé Koekemoer, Carmien Tolmie

## Abstract

*Candida albicans* is a critical priority fungal pathogen causing invasive fungal infections with high mortality rates in immunocompromised patients. The increasing fungal infection rate and resistance of fungal pathogens to existing antifungal treatments have emphasized the need for the development of novel antifungal medicine. The ergosterol biosynthesis pathway has been a successful target for antifungal compounds, but many enzymatic steps remain unexplored. 24C-sterol methyltransferase (24C-SMT) catalyzes a critical fungal-specific step in ergosterol biosynthesis. When 24C-SMT is disrupted, fungal pathogens are sensitized to temperature, various inhibitors, and antifungals, and a loss of virulence can be observed. In this study, five 24C-SMT variants with different lengths of N-termini were heterologously produced in *Escherichia coli* and three were purified to near-homogeneity with immobilized metal-affinity and size-exclusion chromatography. N-terminally truncated *C. albicans* 24C-SMT was utilized for crystallization trials due to its increased stability and higher purity compared to the full-length protein. 24C-SMT crystals were obtained in the presence of Sadenosyl-homocysteine, but diffracted to low resolution. Therefore, we established a starting point for 24C-SMT crystallization by providing an optimized protocol for heterologous 24C-SMT production, purification and initial crystallization conditions, which could be used for further downstream crystallographic studies.

## Introduction

*Candida albicans* is a dimorphic fungus responsible for superficial mucosal and disseminated infections in immunocompromised patients, and topical infections in healthy individuals (Lopes & Lionakis, 2022). Infectious fungal disease, caused by pathogens such as *C. albicans*, has high mortality rates in immunocompromised individuals (up to 88%) and is thus highly relevant in South Africa due to the high per capita HIV/AIDS prevalence (Dos Santos Abrantes et al., 2014). In 2022, the World Health Organization released the first list of priority fungal pathogens, of which *C. albicans, Candida auris, Aspergillus fumigatus* and *Cryptococcus neoformans* are considered critical priority pathogens (World Health Organization, 2022).

Currently, four classes of antifungals are available to treat disseminated fungal infections clinically (Bouz & Doležal, 2021). These are azoles, polyenes, echinocandins and pyrimidines. The widespread use of azoles has led to several fungal species, including non-*albicans Candida* species, developing resistance (Johnson et al., 1995; Pfaller, 2012; Rhodes & Fisher, 2019; Whaley et al., 2017). Polyenes have severe toxic side effects (Dash et al., 2024), and intrinsic resistance to echinocandins has been reported for several pathogens, e.g., *C. neoformans* (Kalem et al., 2021). Furthermore, resistance towards 5-fluorocytosine, the only clinically used pyrimidine, is rapidly developed by *C. albicans* (Lopes & Lionakis, 2022). There is thus a clear and urgent need to develop novel antifungal medicines.

A promising drug target in fungi is ergosterol, a critical sterol that is the counterpart of mammalian cholesterol (Rodrigues, 2018). Ergosterol is regarded as a fungal hormone for its role in stimulating growth and proliferation. In addition, ergosterol is immunoactive and crucial for pathogenesis. Ergosterol and its biosynthesis have been successfully targeted by several antifungal compounds (Bouz & Doležal, 2021). This is evident with the widely used azoles, which target lanosterol 14α-demethylase in the ergosterol biosynthesis pathway, and polyenes, which bind to ergosterol in the fungal cell membrane.

One of the critical enzymes in the ergosterol biosynthesis pathway is 24C-sterol methyltransferase (24C-SMT), encoded by *ERG6* (Nes et al., 2009). 24C-SMT catalyses the transfer of a methyl group from S-adenosyl methionine (SAM) to the C-24 of zymosterol to form fecosterol in yeasts (figure 1). In molds such as *A. fumigatus*, 24C-SMT acts in an alternative pathway by converting lanosterol to eburicol (Alcazar-Fuoli & Mellado, 2013). The conversion of zymosterol by 24C-SMT is the first divergence of ergosterol biosynthesis in fungi from cholesterol biosynthesis in mammals; therefore, the 24C-SMT-catalysed reaction does not occur in mammals and no homologue is present.

**Figure 1:**
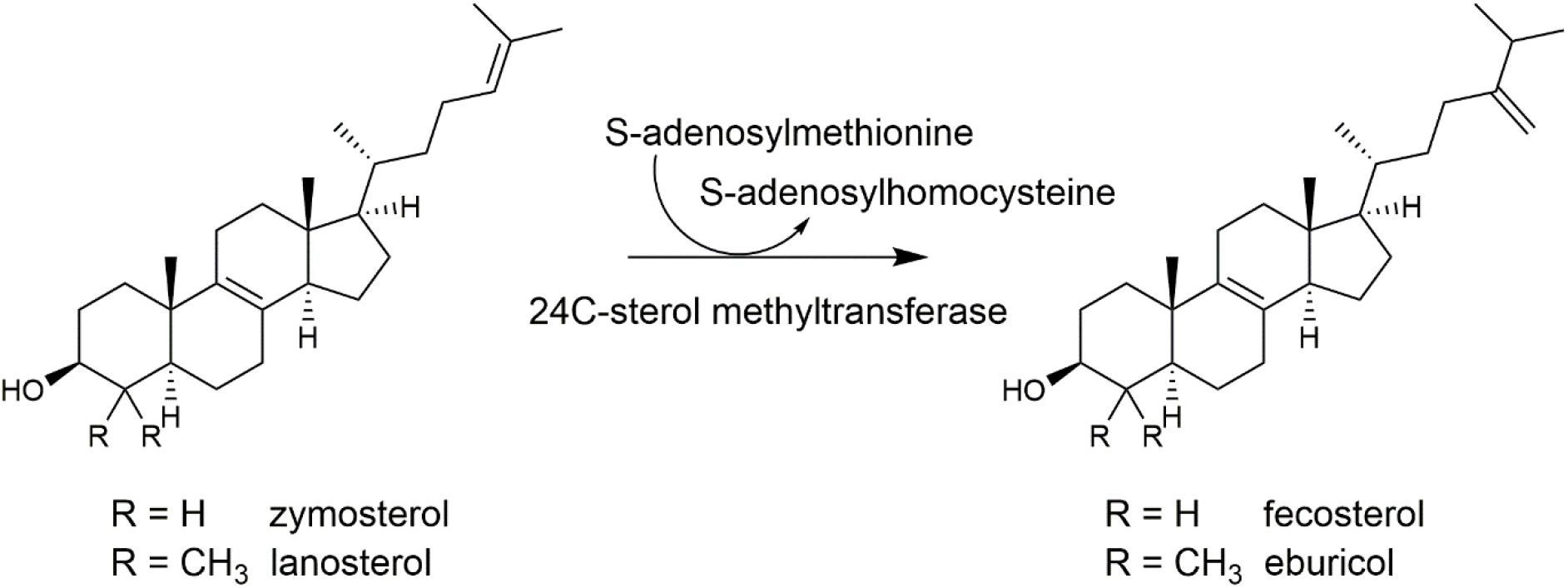
Conversion of zymosterol or lanosterol to fecosterol or eburicol, respectively, by 24C-sterol methyltransferase (24C-SMT, ERG6). Conversion of zymosterol by 24C-SMT typically takes place in yeasts, whereas conversion of lanosterol takes place in molds.

Previous studies have shown that *C. albicans erg6* homozygotes are sensitized to metabolic and ergosterol biosynthesis inhibitors, including terbinafine (Jensen-Pergakes et al., 1998). Similarly, an increased sensitivity towards high osmolarity, salt concentration, increased temperatures and selected antibiotics was observed in *erg6* mutants of *C. neoformans* (Toh et al., 2017). However, the most notable observation is that the echinocandin micafungin inhibited growth in the *erg6* mutants, while the wild-type strain was unaffected. In a more recent study on an *erg6* mutant of *C. neoformans*, loss of 24C-SMT activity inhibited growth at 30°C and 35°C, with growth completely arrested at 37°C (Oliveira et al., 2020). More importantly, *in vivo* experiments with the *erg6* mutant in the invertebrate model organism *Galleria mellonella* (wax moths) showed the mutant strain to be completely avirulent at 37°C. Therefore, 24C-SMT poses an ideal target for developing novel antifungal compounds.

To date, no crystal structure has been reported for any 24C-SMT in the Protein Data Bank (PDB). Structure predictions of *C. albicans* 24C-SMT with AlphaFold2 (Jumper et al., 2021) showed a well-folded globular protein, and bioinformatics analysis indicated that no extensive intrinsically disordered regions were detected. Therefore, we anticipated that this enzyme would be amenable to crystallization. Here, we report the expression, purification, and crystallization of truncation libraries of 24C-SMT from the pathogenic fungus *C. albicans* as a novel antifungal drug target for structure-based drug discovery studies.

## Materials and Methods

### In silico construct design

A model of the full-length *C. albicans* 24C-SMT (UniProt accession no. O74198.2) was generated using the SWISS-MODEL server <u>SWISS-MODEL (expasy.org)</u> (Bienert et al., 2017; Guex et al., 2009; Waterhouse et al., 2018). The model was inserted into ESPript/ENDscript (<u>ESPript 3.x / ENDscript 2.x (ibcp.fr)</u> (Robert & Gouet, 2014) to obtain a prediction of the secondary structure, and to provide a structural alignment with *Saccharopolyspora spinosa* [4+2]-cyclase SpnF (PDB ID: 4PNE) (Fage et al., 2015). The alignment revealed that 4PNE had no modeled structure until the equivalent residue of Leu66 in *C. albicans* 24C-SMT. The 24C-SMT protein was then truncated on the N-terminus on a helix-by-helix approach. 24C-SMT variant 1 is the full-length protein. In variants 2 – 6, the first 9, 28, 37, 55 and 67 residues were removed, respectively, corresponding to the truncation of no α-helices but a potentially unstructured region, α-helix 1, α-helices 1-2, α-helices 1-3 and α-helices 1-4 (figure 2).

**Figure 2:**
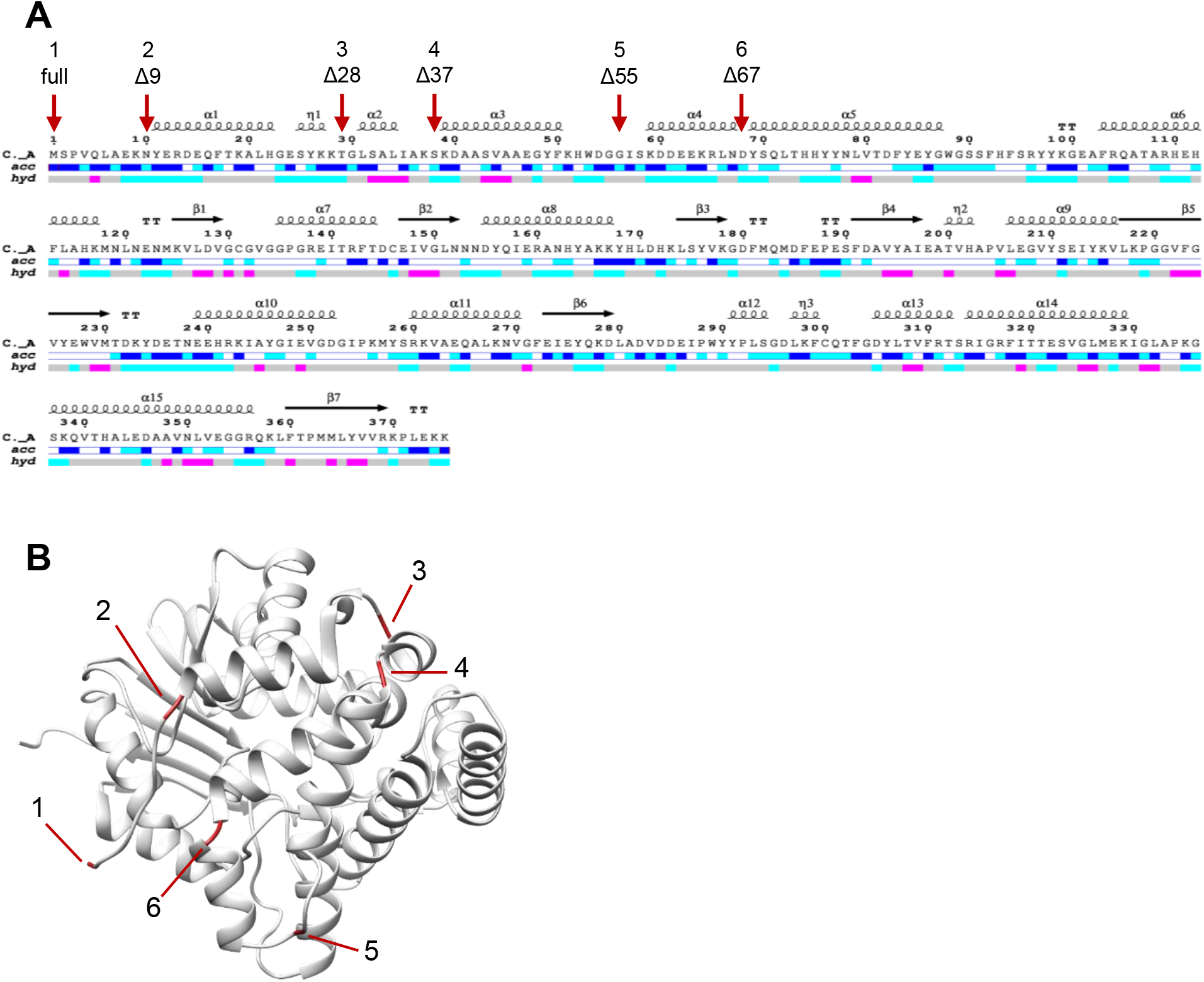
Truncations of the N-terminus of 24C-SMT (variants 1-6). Five truncations were designed on a helix-by-helix approach. **A**. The N-termini of truncated variants are shown by the red arrows on the EndScript output, and the number of residues removed is indicated by Δ. **B**. The truncations as shown on the protein model (red indicates the N-terminus of the variant).

### PCR amplification of C. albicans ERG6 truncations

The *C. albicans ERG6* gene was codon-optimized for *Escherichia coli* and commercially synthesized (Twist Bioscience) and received in the pET-28a(+) plasmid. Truncations of the *ERG6* gene were amplified from pET-28a(+):*ERG6* with forward primers containing the 5’ Golden Gate extension (5’-AAAAAA<u>GGTCTC</u>ACATG-target-3’, *Bsa*I restriction site underlined), where ATG is the start codon of the target *ERG6* gene. The reverse primers were similarly designed, but the 5’ extension was 3’-target-TAATGACTCGA<u>GAGACC</u>AAAAAA-5’, with the complement 5’-TTTTTTGGTCTCTCGAGTCATTA-tegrat-3’. For the vectors encoding a C-terminal 6xHis-tag, the stop codons (TAATGA) were removed from the primers, whereas the remainder of the sequences were the same. The primers used to amplify the *ERG6* full-length gene and truncations with their respective melting temperatures (T_m_) are listed in Supplementary table 1.

The Q5 High-Fidelity DNA Polymerase (New England Biolabs) was used for amplification, and the PCR reaction consisted of 1X Q5 reaction buffer, 200 μM dNTPs, 0.5 μM forward primer, 0.5 μM of reverse primer, 1 μL template DNA, 1 U Q5 DNA polymerase and nuclease-free water to a final volume of 25 μL. The T100 thermocycler (BioRad) with a heated lid (105°C) was used to run the amplification protocol. Thermocycling began with initial denaturation at 98°C for 30 seconds, followed by 34 cycles of 98°C for 10 seconds, annealing of the primers to the template for 20 seconds at the primer T_m_ and extension at 72°C for 30 seconds. These steps were repeated for 34 cycles before a final extension step at 72°C for 2 minutes. The size of the PCR product was determined on a 1% (w/v) agarose gel. The PCR product was cleaned up using the QIAquick PCR purification kit (Qiagen), and the DNA concentration was determined using a NanoDrop spectrophotometer (Thermo Fisher Scientific).

A *Dpn*I digestion was performed post-PCR to remove the methylated template DNA as it has the same antibiotic resistance (kanamycin) as the destination Golden Gate vector. This reaction consisted of 25 μL of PCR product, 20 U *Dpn*I restriction enzyme (New England Biolabs) and 1X rCutSmart Buffer. The reaction was incubated at 37°C for 30 minutes and incubated further at 80°C for 20 minutes to inactivate the *Dpn*I enzyme.

### Golden Gate cloning assembly reaction

The Golden Gate cloning reaction was prepared by adding 100 ng of either pNIC-CTH10STII (encoding a C-terminal 6xHis-tag) or pNIC-NHSTIIT-GG vector (encoding an N-terminal 6xHis-tag) (Fairhead et al., 2024) with a 3-fold molar excess of the purified PCR product (NEBioCalculator, https://nebiocalculator.neb.com/#!/ligation). To complete the reaction, 1X ligase buffer, 5 U T4 DNA ligase, 10 U *Bsa*I-HFv2, and deionized water were added to a final volume of 10 μL (New England Biolabs).

The T100 thermal cycler with a heated lid (105°) was used for the thermocycling process. The reaction was initiated with digestion of the vector and insert at 37°C for 5 minutes, followed by ligation of the DNA fragments at 16°C for 5 minutes. These steps were repeated for 10 cycles. This was followed by digestion of the remaining unmodified vector at 37°C for 10 minutes. Thereafter, the reaction was deactivated at 80°C for 20 minutes.

The ligation reaction was transformed into CCMB80 (Von der Haar, 2019) Mach1 *E. coli* competent cells (Thermo Fisher Scientific). After selection on LB-kanamycin plates (50 μg/mL), a colony PCR reaction was used to confirm whether the insert was cloned into the vector, consisting of 5 U MyTaq DNA polymerase (Bioline), 1X MyTaq reaction buffer, and 0.4 μM T7 promoter and T7 terminator primers. The T100 thermal cycler with a heated lid (105°) was used for the thermocycling process. Thermocycling began with a denaturation step at 95°C for 1 minute, followed by a denaturation step at 95°C for 15 seconds, an annealing step at 55°C for 15 seconds, and an extension step at 72°C for 60 seconds, which was repeated for 35 cycles. This was followed by a final elongation step at 72°C for 1 minute. The PCR products were run on a 1% (w/v) agarose gel to determine the presence of the correctly sized insert. Sanger sequencing was conducted on the positive clones with the T7 promoter and T7 terminator primers to confirm successful cloning.

### Test expression and purification

*Escherichia coli* BL21(DE3) cells (Centre for Medicines Discovery, University of Oxford) were transformed with 100 ng of plasmid, and a single colony was inoculated into 1 mL of AIM-TB (ForMedium) supplemented with kanamycin (50 μg/mL) in a 24-well plate. Cells were initially incubated for 4 hours at 37°C and further at room temperature overnight, at 250 rpm. Cells were lysed and 24C-SMT purified using the MagneHis protein purification system (Promega) as outlined in the manufacturer’s protocol. The presence of the His-tagged protein was determined with SDS-PAGE using NuPAGE 4-12% Bis-Tris Midi Protein Gels (Thermo Fisher Scientific).

Upscaling for purification was performed as previously described in the PREPX workflow (Fairhead, 2024). Briefly, freshly transformed *E. coli* BL21(DE3) cells were collectively streaked and inoculated into 10 mL LB media containing 50 μg/mL kanamycin and 0.5% (w/v) glucose. The starter culture was grown for 4 hours at 37°C and 250 rpm shaking before being inoculated into 1 L of AIM-TB, supplemented with 50 μg/mL kanamycin and 0.01% (v/v) Antifoam-204 (Merck), in a 2.5 L baffled shake flask. Cells were incubated for 4 hours at 37°C and then overnight at 17°C and 200 rpm. Cells were harvested by centrifugation at 4000xg for 20 minutes at 4°C, and the pellet (cell weight ∼50 g/L) was stored at -80°C.

For cell lysis, the pellets were re-suspended using 3 mL of Base Buffer (10 mM HEPES, 500 mM NaCl, 5% glycerol (v/v), 30 mM imidazole, 0.5 mM TCEP, pH 7.5) per gram of cell weight. Triton X-100 (1% (v/v)), lysozyme (0.5 mg/mL) and benzonase (1 μg/mL) were also added and the pellet was incubated for 30 minutes at room temperature before freezing at -80°C. The cell lysate mixture was thawed in a room-temperature water bath and then centrifuged at 4800xg for 1 hour at 4°C. The soluble fraction was loaded onto a His GraviTrap column (Cytiva) equilibrated with binding buffer (Base Buffer containing 30 mM imidazole). The columns were washed with 2 x 10 mL of binding buffer prior to eluting the His-tagged target protein with 2.5 mL of Base Buffer containing 500 mM imidazole. The protein was immediately desalted with a PD-10 column (Cytiva) using binding buffer. The protein concentration was determined using absorbance at 280 nm on a NanoDrop spectrophotometer. The His-tag was removed by protease digestion overnight at 4°C using 1 mg of TEV protease (Kapust et al., 2001) per 10 mg of target protein. The tag, uncleaved protein and TEV protease (bearing a non-cleavable His-tag) were then removed by passing through a His GraviTrap column, using binding buffer. The protein was concentrated using an Amicon Ultra centrifugal concentrator (30 kDa MWCO, Merck) before being loaded on an equilibrated Superose 12 pg column (Cytiva) for size-exclusion chromatography using Base Buffer. The efficacy of the process and final protein purity were assessed by SDS-PAGE, and the protein was stored at -80°C.

### Crystallization

Crystallization was performed using the sitting-drop vapor diffusion method with the mosquito crystallization robot (SPT Labtech) in SWISSCI 96-well 3 Lens Crystallization plates at 20°C using a Rock Imager (Formulatrix). The first images were taken after 12 hours, and the imaging schedule followed a Fibonacci sequence of days for further collections. Coarse screens for initial crystallization trials were set up with 23 mg/mL protein and SAM (2 mM) or S-adenosyl-homocysteine (SAH, 1.3 mM). Coarse screens tested included the Ligand-friendly screen, ShotGun1, BCS, customized version of JCSG plus, customized version of Hampton Crystal screen, customized version of Hampton Index screen (all from Molecular Dimensions). Follow-up screens were designed and dispensed using the Formulator Screen Builder for Protein Crystallography (Formulatrix). Crystals were cryoprotected using 33% (v/v) ethylene glycol, glycerol or PEG300 and cryocooled. X-ray diffraction data was collected using UDC mode at beamline I03, Diamond Light Source, UK, as part of the CMD BAG allocation mx34598.

## Results

### Cloning

The full-length *C. albicans ERG6* gene and N-terminally truncated variants were PCR-amplified from the pET-28a(+):*ERG6* construct (figure 3) and subcloned into the pNIC-CTH10STII and pNIC-NHSTIIT-GG vectors. Colony PCR was used to determine the presence or absence of the *ERG6* gene insert (figure 4). Colony PCR screening showed that all *ERG6* genes were successfully subcloned into their respective vectors, with the exception of variant 3 in pNIC-NHSTIIT-GG. Positive clones were further verified with Sanger sequencing.

**Figure 3:**
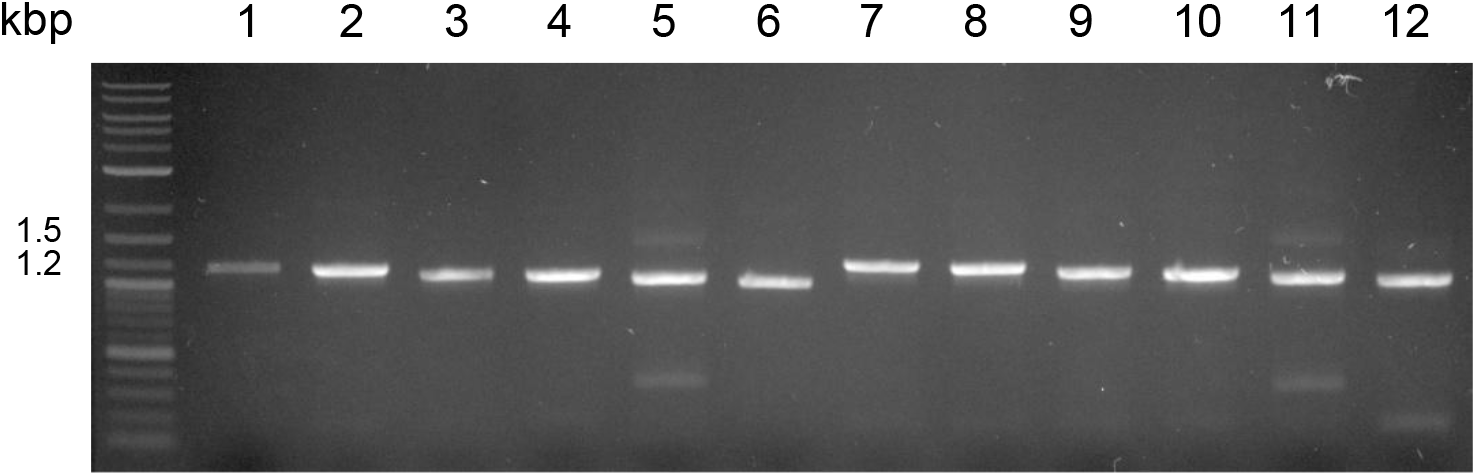
Agarose gel electrophoresis of *C. albicans ERG6* PCR amplicons (∼1200 bp). Wells 1-6, variant 1-6 amplicons (incl. stop codon) to be cloned into the pNIC-NHSTIIT-GG vector encoding an N-terminal His-tag. Wells 7-12, variant 1-6 amplicons (excl. stop codon) to be cloned into the pNIC-CTH10STII-GG vector encoding a C-terminal His-tag.

**Figure 4:**
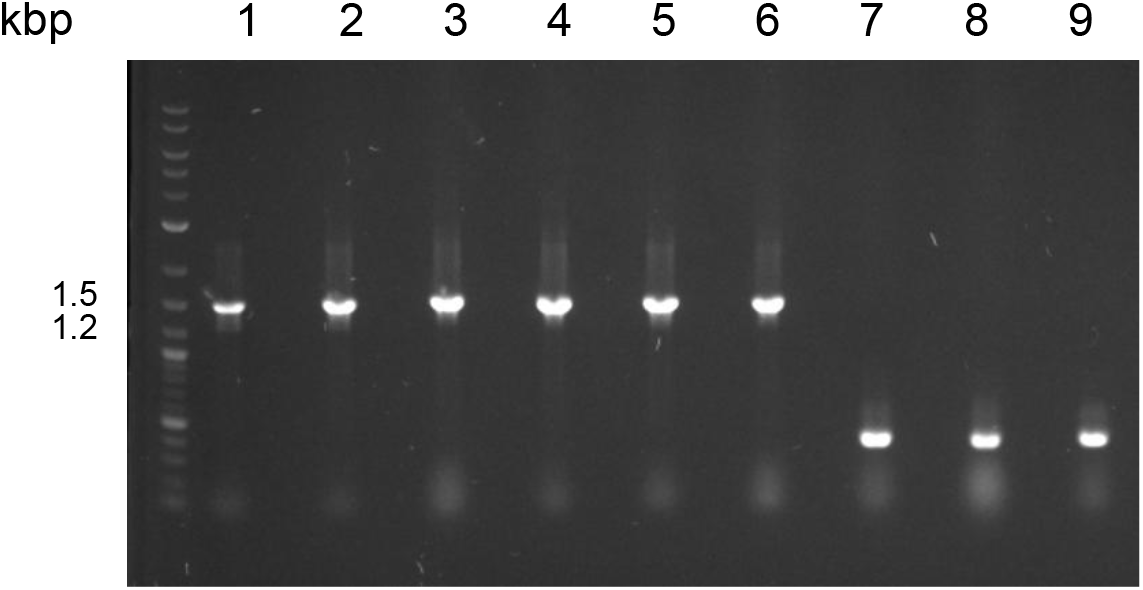
Agarose gel electrophoresis of colony PCR for *C. albicans ERG6* variants in pNIC-NHSTIIT. Wells 1-3, *ERG6* variant 1; wells 4-6, variant 2; wells 7-9, variant 3. *ERG6* variants 1 and 2 represented the successful clones (wells 1-6 containing the insert of ∼1500 bp) and *ERG6* variant 3 were failed clones (wells 7-9). In total, 11/12 *ERG6* constructs were successfully cloned.

### Protein expression and purification

A test expression in *E. coli* and a small-scale purification using Ni-NTA magnetic beads were conducted for the 11 N- or C-terminally His-tagged *C. albicans ERG6* constructs. SDS-PAGE analysis indicated that the *C. albicans* pNIC-NHSTIIT-*ERG6* and pNIC-CTH10STII-*ERG6* constructs were expressed, with the exception of the full-length 24C-SMT with a C-terminal His-tag, which had very low or no expression. The 24C-SMT proteins bound to and eluted from the Ni-NTA beads (figure 5, ∼40kDa). However, expression yields varied between constructs, with the pNIC-NHSTIIT-*ERG6* truncation variants 4, 5 and 6 having the highest expression yields. Therefore, these three variants were selected to upscale for purification. Although the pNIC-CTH10STII-*ERG6* constructs showed production of the 24C-SMT protein, purification was unsuccessful as the protein precipitated out of solution during IMAC and ultimately, could not be eluted. Hence, no C-terminally His-tagged protein was obtained.

**Figure 5:**
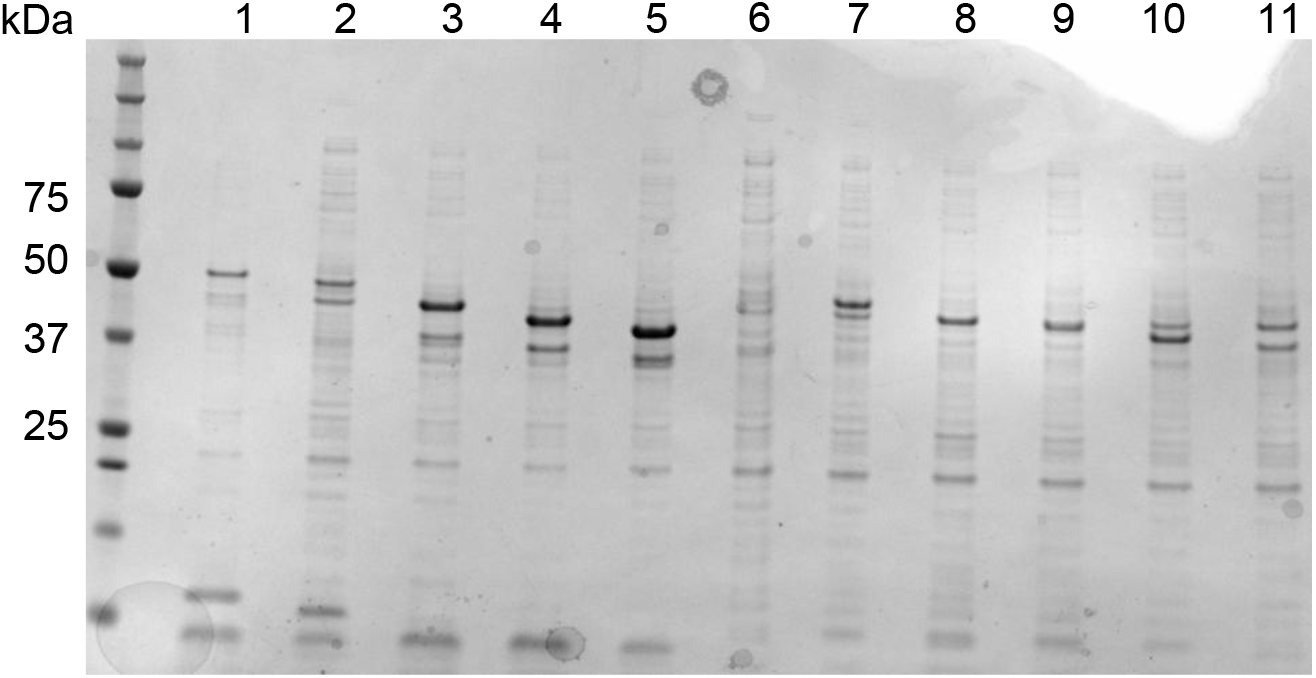
SDS-PAGE analysis of test expressions of *C. albicans* pNIC-NHSTIIT-*ERG6* and pNIC-CTH10STII-*ERG6* constructs. Wells 1-5, pNIC-NHSTIIT-*ERG6* variants 1, 2, 4, 5 and 6, respectively (49-42 kDa). Wells 6-11, pNIC-CTH10STII-*ERG6* variants 1-6, respectively (48-41 kDa).

Variants 4, 5 and 6 were purified with IMAC, and the His-tag was successfully cleaved using TEV protease and removed with reverse IMAC (figure 6). The tag-free proteins were further purified with size-exclusion chromatography, which yielded proteins corresponding to the predicted sizes for all three truncation variants (variant 4: 39.1 kDa, variant 5: 37.2 kDa, variant 6: 35.8 kDa). Notably, proteolysis was still evident after size-exclusion chromatography in variant 4, where the sequence encoding the N-terminal first and second α-helices (α1-2) were removed, and in variant 6, where the α1-4 helices were removed. However, variant 5, lacking the α1-3 helices, remained intact; thus, this variant was used for crystallization trials.

**Figure 6:**
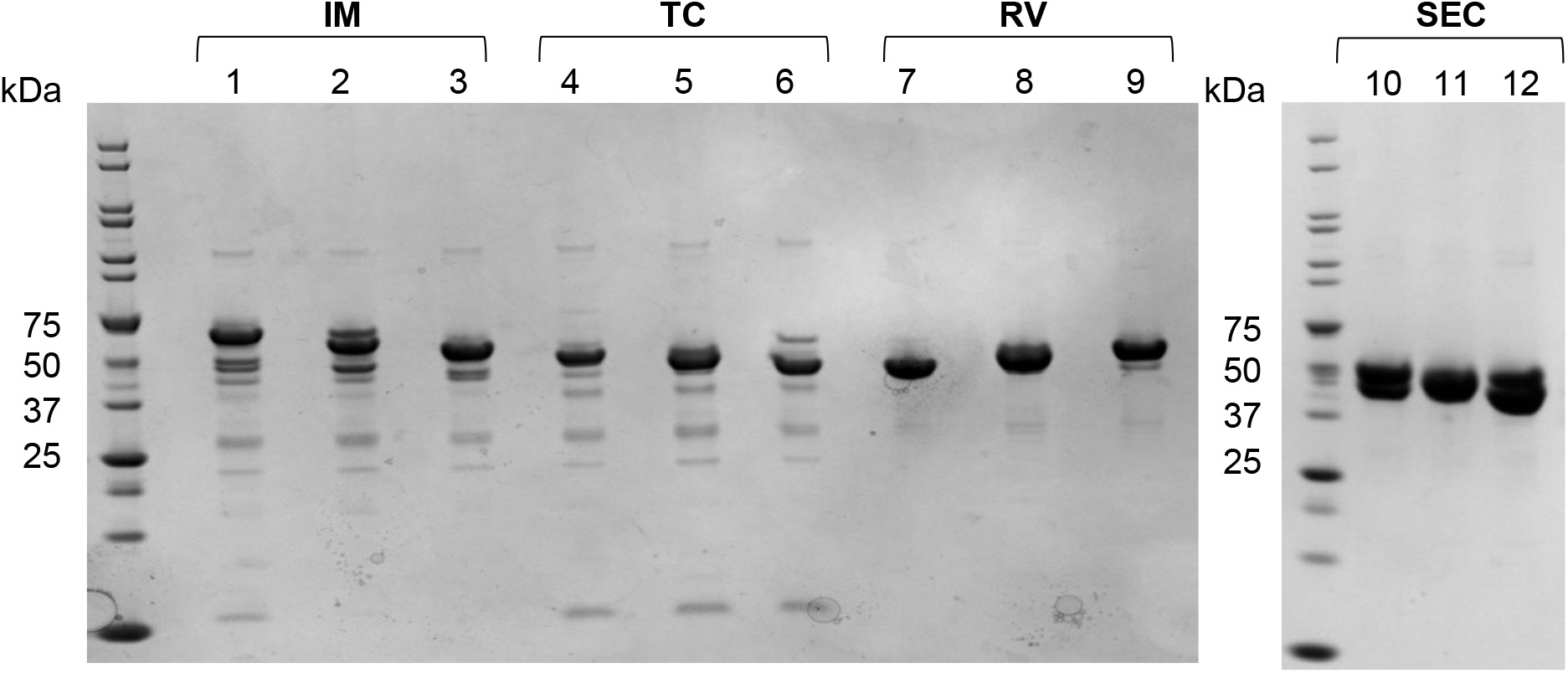
SDS-PAGE analysis of the purification of *C. albicans* 24C-SMT truncation variants 4, 5 and 6. Wells 1-3, His GraviTrap eluate (IM), where well 1-variant 4, well 2-variant 5 and well 3-variant 6 (44-41 kDa). Wells 4-6, protein after overnight TEV protease cleavage (TC), where well 4-variant 4, well 5-variant 5 and well 6-variant 6. Wells 7-9, reverse IMAC eluate after tag cleavage (RV), where well 7-variant 6, well 8-variant 5 and well 9-variant 4 (35-39 kDa). Wells 10-12, variants after size-exclusion chromatography (SEC) with well 10-variant 4 (39 kDa), well 11-variant 5 (37 kDa) and well 12-variant 6 (36 kDa).

### Crystallization and diffraction data collection

Crystals were obtained after four days using the sitting-drop vapor-diffusion method at 20°C (figure 7) in the condition containing 1.3 mM SAH, 0.1 M tri-sodium citrate, 17% PEG Smear Broad and 0.15 M magnesium acetate. No crystals were obtained in the apo form, and co-crystallization attempts with SAM resulted in immediate precipitation of the protein. Unfortunately, the crystals only diffracted to ∼5-7 Å.

**Figure 7:**
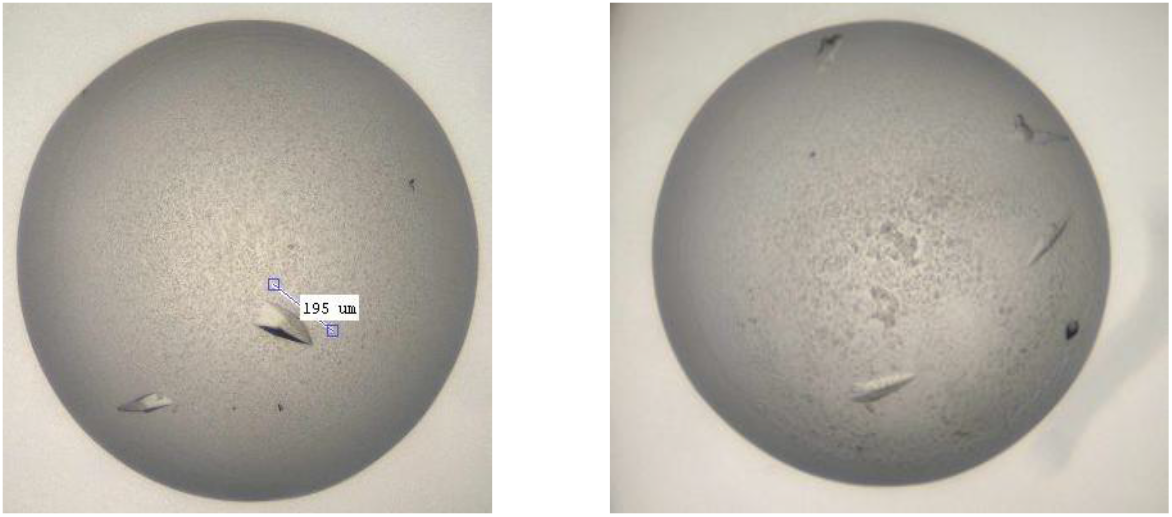
*C. albicans* SMT variant 5 crystals co-crystallized with S-adenosyl homocysteine.

## Discussion and Conclusions

X-ray crystallographic fragment screening has been used to develop inhibitor compounds against a wide range of targets from diverse viruses (Bauman et al., 2013; Munawar et al., 2018; Newman et al., 2021; Patel et al., 2016), bacteria (Füsser et al., 2023; Garbers et al., 2024; Willocx et al., 2025) and parasites (de Souza Neto et al., 2024). However, to our knowledge, this technique has not yet been exploited in the development of antifungal compounds. This study aims to provide an optimized experimental protocol to heterologously produce and purify high-quality *C. albicans* 24C-SMT for downstream crystallographic structural studies and fragment screening experiments.

Previously, purification of the full-length *C. albicans* 24C-SMT produced in *E. coli* yielded protein with multiple truncated forms (unpublished, Supplementary figure S1). Altering the media composition, incubation time and temperature did not remedy this, and while some improvement was seen by co-expressing with the GroES/EL chaperone, truncated forms were still present. Hence, we turned to producing truncated variants of the protein.

In this study, 11 construct variants of the *C. albicans* 24C-SMT were created, encoding cleavable N-terminal (5 variants) or C-terminal (6 variants) 6xHis-tags, by subcloning into the pNIC-NHSTIIT and pNIC-CTH10STII vectors, respectively. N-terminal truncation of at least the first two predicted α-helices, combined with an N-terminal His-tag, were effective in improving the overall heterologous expression and purification of 24C-SMT. Tag cleavage with TEV protease, followed by reverse IMAC to remove the cleaved tag, uncleaved protein and other contaminants, significantly improved the purity of the protein.

Notably, over- or undertruncation increased the susceptibility to proteolysis, resulting in a non-homogeneous population of purified protein. Ultimately, only one variant remained intact and thus suitable for crystallization. Unfortunately, the crystals obtained only diffracted to very low resolution. Nevertheless, these initial crystals confirm that the 24C-SMT variant is crystallizable and provides an avenue for optimization of the crystallization conditions, possibly using microseeding (D’Arcy et al., 2007), to improve the diffraction resolution.

## Acknowledgements

This work is based on the research supported in part by the National Research Foundation of South Africa (TTK23041793934 and PMSD230511104743), the Africa Oxford Initiative (Catalyst Grant AfOx-264), the University of the Free State (South Africa), and the Agricultural Research Foundation (South Africa). The authors would like to thank the Protein crystallography small research facility (PX-SRF, CMD, University of Oxford) for hosting the research exchange essential to the success of this work, as well as the beamline scientists of the Diamond Light Source beamline I03. Diffraction data was collected under proposal mx34598.

## Supplementary data

**Table 1.**
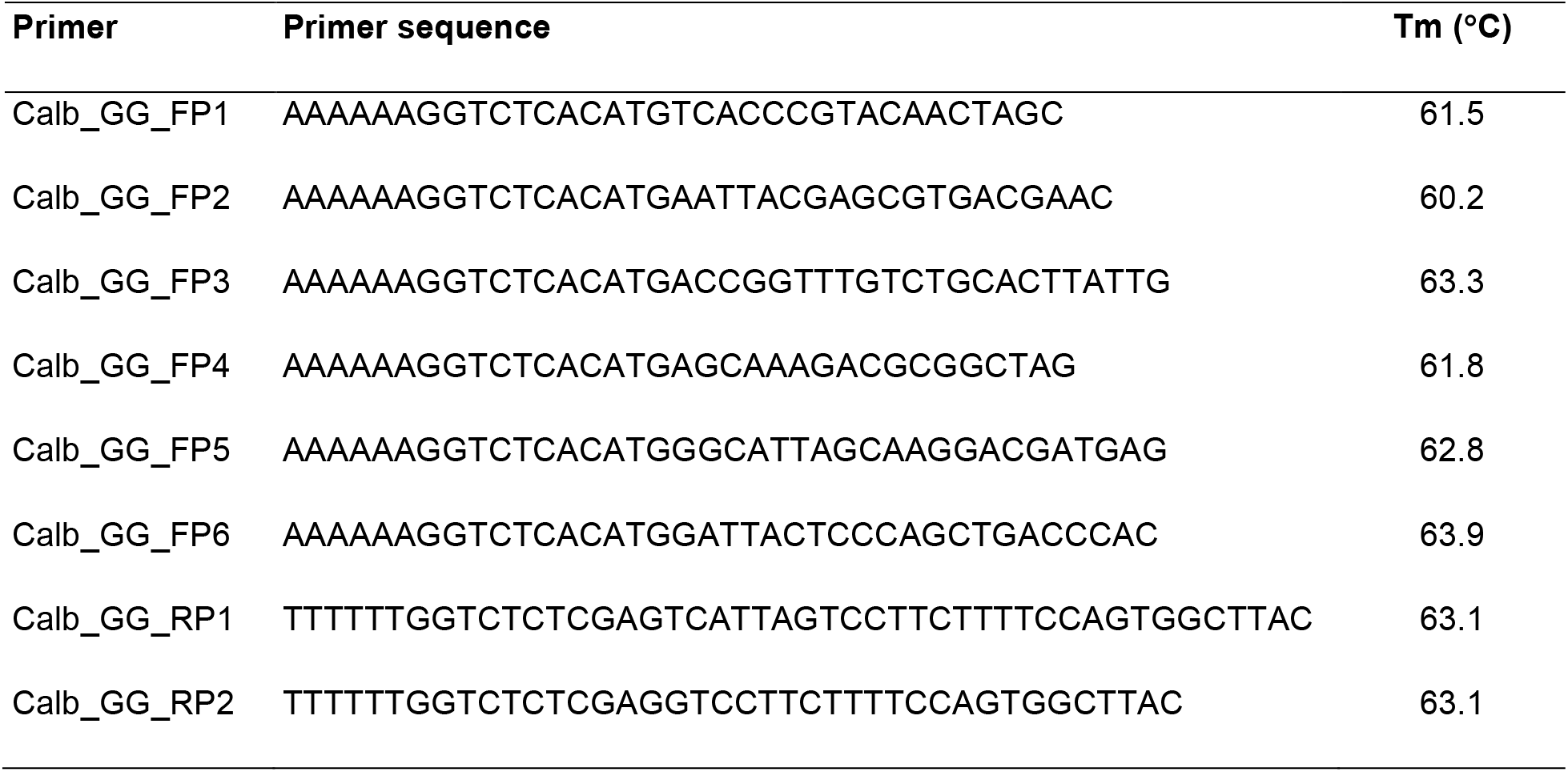
Primer sequences used to amplify *ERG6* truncation variants.

**Supplementary sequence 1:** Amino acid sequence of pNIC-NHSTIIT-GG-encoded ERG6 construct variant 1. * denotes TEV protease cleavage site. The original start codon (residue 1) indicated by <u>M</u>. Calculated MW= 48852.618 Da before tag removal and 43158.63 Da after tag removal, (ε)=73230 M^-1^cm^-1^ before tag removal and 70375 M^-1^cm^-1^ after tag removal.

MHHHHHHSSGASWSHPQFEKGGGSGGGSGGSAWSHPQFEKGSGVDLGTENLYFQ*SM SPVQLAEKNYERDEQFTKALHGESYKKTGLSALIAKSKDAASVAAEGYFKHWDGGISKDDEEKRLNDYSQLTHHYYNLVTDFYEYGWGSSFHFSRYYKGEAFRQATARHEHFLAHKMNLNENMKVLDVGCGVGGPGREITRFTDCEIVGLNNNDYQIERANHYAKKYHLDHKLSYVKGDFMQMDFEPESFDAVYAIEATVHAPVLEGVYSEIYKVLKPGGVFGVYEWVMTDKYDETNEEHRKIAYGIEVGDGIPKMYSRKVAEQALKNVGFEIEYQKDLADVDDEIPWYYPLSGDLKFCQTFGDYLTVFRTSRIGRFITTESVGLMEKIGLAPKGSKQVTHALEDAA VNLVEGGRQKLFTPMMLYVVRKPLEKKD

