## Supplementary figures and images for "Heterologous production, purification and crystallization of 24C-sterol methyltransferase from *Candida albicans*"

### Supplementary figure 1

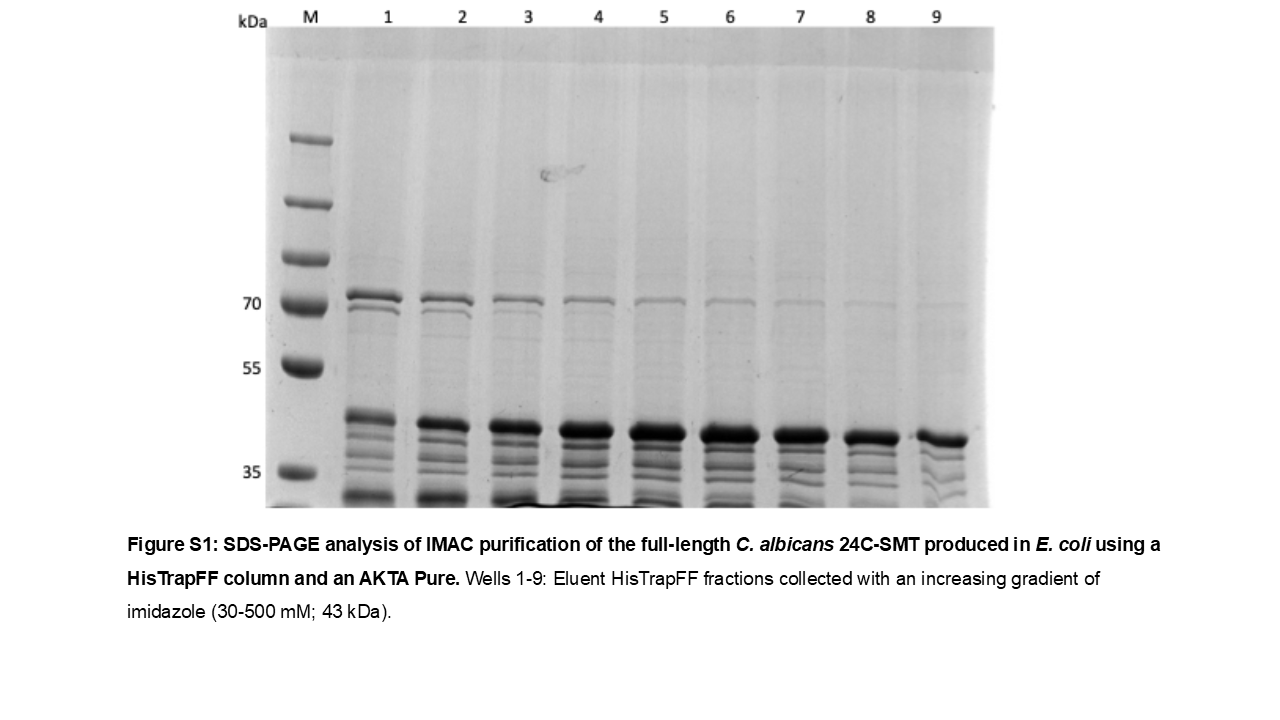
